# Remodeling Pathway Control of Oxidative Phosphorylation by Temperature in the Heart

**DOI:** 10.1101/103457

**Authors:** Hélène Lemieux, Pierre U. Blier, Erich Gnaiger

## Abstract

The capacity of mitochondrial oxidative phosphorylation (OXPHOS) and fuel substrate supply are key determinants of cardiac muscle performance. Although temperature exerts a strong effect on energy metabolism, until recently numerous respiratory studies of mammalian mitochondria have been carried out below physiological temperature, with substrates supporting submaximal respiratory capacity. We measured mitochondrial respiration as a function of temperature in permeabilized fibers from the left ventricle of the mouse heart. At 37 °C, OXPHOS capacity with electron entry through either Complex I or Complex II into the Q-junction was about half of respiratory capacity with the corresponding physiological substrate combination reconstituting tricarboxylic acid cycle function with convergent electron flow through the NADH&succinate (NS) pathway. When separating the component core mitochondrial pathways, the relative contribution of the NADH pathway increased with a decrease of temperature from 37 to 25 ºC. The additive effect of convergent electron flow has profound consequences for optimization of mitochondrial respiratory control. The apparent excess capacity of cytochrome *c* oxidase (CIV) was 0.7 above convergent NS-pathway capacity, but would be overestimated nearly 2-fold with respect to respiration restricted by provision of NADH-linked substrates only. The apparent excess capacity of CIV increased sharply at 4 °C, caused by a strong temperature dependence of and OXPHOS limitation by NADH-linked dehydrogenases. This mechanism of mitochondrial respiratory control in the hypothermic mammalian heart is comparable to the pattern in ectotherm species, pointing towards NADH-linked mt-matrix dehydrogenases and the phosphorylation system rather than electron transfer complexes as the primary drivers of thermal sensitivity at low temperature and likely modulators of temperature adaptation and acclimatization. Delineating the link between stress and remodeling of OXPHOS is critically important for improving our understanding of metabolic perturbations in disease evolution and cardiac protection. Temperature is not a trivial experimental parameter to consider when addressing these questions.

## Introduction

Contractile activity in cardiac muscle mainly depends on mitochondrial (mt) energy generated by oxidative phosphorylation (OXPHOS). The heart is highly sensitive to defects in OXPHOS [1], stress-induced mitochondrial cytopathies and degenerative mitochondrial defects, including heart failure [2], acute ischemia and myocardial infarct [3], ischemia-reperfusion [4], type 2 diabetes [5–7], aging [8, 9], and inherited genetic diseases [10–12]. Patients present with functional impairment when the capacity of an enzyme is reduced below its threshold activity. This threshold effect is a function of the apparent excess enzyme activity above pathway capacity. To evaluate the threshold and excess capacity of a single step in OXPHOS, it is necessary not only to quantify the changes in enzyme activity, but determine the impact of these changes on respiratory capacity through mt-pathways representative of physiological conditions [13].

Respiration in the mammalian heart is supported by carbohydrates (10 to 40% [14]) and fatty acids (60 to 90% [15]). Electron transfer in the NADH- and succinate-linked pathway (NS) converges through Complex I and Complex II at the Q-junction [16] (Fig 1A). Downstream electron flow is catalyzed by Complex III and Complex IV (cytochrome *c* oxidase) to oxygen as the terminal electron acceptor. Conventional protocols in bioenergetics use either NADH-linked substrates (N) or succinate&rotenone (S), thereby separating the system into linear thermodynamic cascades, forming distinct electron transfer chains [17, 18]. This experimental design aims at the measurement of biochemical coupling efficiency and proton stoichiometry, and is applied in the functional diagnosis of specific OXPHOS defects. As recognized in mitochondrial physiology, however, it does not allow estimation of maximal respiratory capacity under physiological conditions. Fuel substrates supporting convergent electron transfer at the Q-junction enhance respiratory capacity, as shown when succinate is added to NADH-linked substrates, reconstituting physiological tricarboxylic acid cycle function with combined NS-pathway flux. This effect of succinate varies depending on species, strains, organ and experimental conditions; stimulation is 1.6 to 2.0-fold in rat heart [19, 20], 1.2 to 1.8-fold in rat skeletal muscle [21–23], 1.4-fold in mouse skeletal muscle [24], and 1.3 to 2.1-fold in human skeletal muscle (reviewed by Gnaiger [25]). Similarly, glycerol-3-phosphate (Fig 1A) exerts an additive effect on respiration when combined with pyruvate&malate in rabbit skeletal muscle mitochondria [26], and stimulates respiration above the level of NS-linked oxygen consumption in human lymphocytes [27]. Such substrate combinations do not exert completely additive effects on flux due to intersubstrate competition for transport across the inner mt-membrane [28] and flux control by limited enzyme capacities downstream of the Q-junction.

**Fig1.**
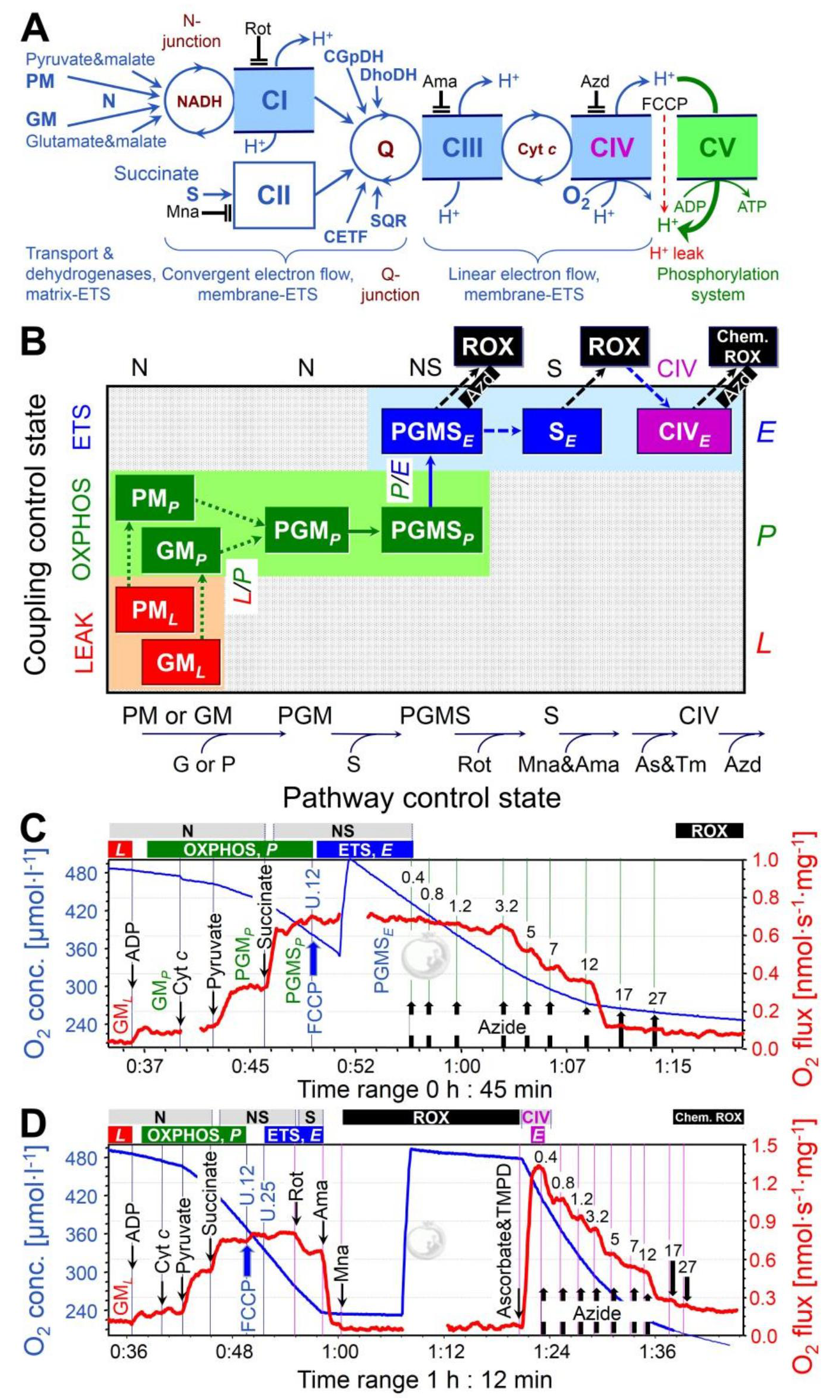
Mitochondrialpathways,substrate-uncoupler-inhibitor-titration(SUIT)protocols and respiration of permeabilized cardiac fibers. **(A)** Schematic representation of the electron transfer system(ETS) coupled to the phosphorylation system (ATP synthase, adenylate translocator and inorganic phosphate transporter). Electron flow from pyruvate&malate; (PM) or glutamate&malate; (GM) converges at the N-junction (NADH-cycle). Electrons converge at the Q-junction from Complex I (CI, NADH-ubiquinone oxidoreductase), Complex II (CII, succinate-ubiquinone oxidoreductase), glycerophosphate dehydrogenase Complex (CGpDH), electron-transferring flavoprotein Complex (CETF), dihydro-orotate dehydrogenase (DhoDH [29]), sulfide-ubiquinone oxidoreductase (SQR [30]), and choline dehydrogenase (not shown), followed by a linear downstream segment through Complexes III (CIII, ubiquinol-cytochrome *c* oxidoreductase) and CIV (cytochrome *c* oxidase), to the final electron acceptor oxygen. CI, CIII, and CIVare proton pumps generating an electrochemical potential difference across the inner mt-membrane. Coupling of the phosphorylation system with the ETS allows the proton potential to drive phosphorylation of ADP to ATP (coupled flow). Protonophores such as FCCP uncouple the ETS from ATP production. Rotenone, malonate and antimycin A are specific inhibitors of CI, CII and CIII, respectively, and were sequentially added at saturating concentrations. **(B)** Coupling/pathway control diagram illustrating the two protocols starting with either PM or GM (SUIT 1 and 2), convergent electron flow at the Q-junction in the NADH&succinate (NS) pathway, and azide titrations in the NS-pathway control state or single step of CIV. **(C)** SUIT 2a with azide titration in the NS-pathway control state. **(D)** SUIT 2b with azide titration in the CIV single enzyme step as a basis of threshold plots.

The aim of the present study is to quantify respiratory capacity of mouse heart mitochondria when providing reducing equivalents simultaneously from the N-pathway (feeding electrons into CI) and succinate (S, feeding electrons into CII) converging at the Q-junction. We determined the impact of temperature on respiratory capacity under these conditions. With NADH-linked substrates, the major flux control resides upstream in the dehydrogenases of the citric acid cycle, with a high apparent excess capacity of respiratory complexes downstream [13]. In the physiological state of combined NS-electron supply, flux control is shifted downstream. We therefore examined the apparent excess capacity of CIV over NS-linked convergent pathway flux, and determined the limitation of NS-OXPHOS capacity by the phosphorylation system (Fig 1A). Mammalian mitochondria are frequently studied at 25 or 30 °C [31]. Temperature coefficients are used to extrapolate the results to 37 °C, but are available for only a few metabolic states [32]. We therefore determined respiratory flux in a variety of mitochondrial coupling/pathway control states at 25, 30 and 37 °C, and extended our study to hypothermia (cold storage temperature, 4 °C) and hyperthermia (40 °C). Modest and deep hypothermia reduce myocardial metabolism temporarily and reversibly, and limit ischemic damage during cardiac surgery [33–37], organ preservation [38, 39], and preservation of mitochondrial function during preparation of isolated mitochondria and permeabilized fibers [40].

Understanding the control of cell respiration and health and disease requires (i) measurement of mitochondrial pathway control at physiologically and clinically relevant temperatures and (ii) application of substrate combinations which are representative of cellular conditions. Our results clearly show that those two criteria dramatically change our estimation of maximal respiratory capacity and control of mitochondrial respiration, providing a reference for physiological and pharmacological intervention strategies. Our results also provide new perspectives on evolutionary temperature adaptation, suggesting that key control of energy transformation and survival at low temperatures might not primarily be exerted by electron transfer complexes but by (i) upstream elements of the ETS including transport of substrates across the inner mitochondrial membrane and matrix dehydrogenases, and (ii) downstream elements of the phosphorylation system.

## Results

### Pathway Control of Mitochondrial Respiratory Capacity

We quantified mitochondrial respiratory capacity in substrate-uncoupler-inhibitor-titration (SUIT) protocols (Fig 1 and 2). Fuel substrates of the N-pathway reduce NAD^+^ to NADH at five key steps: pyruvate dehydrogenase, isocitrate dehydrogenase, glutamate dehydrogenase, oxoglutarate dehydrogenase and malate dehydrogenase. Addition of pyruvate (P) to glutamate&malate (GM) increased OXPHOS capacity (PGM_*P*_) by a factor of 2.5 (2.0-3.7) at 37 ºC (Fig 2A, B). In contrast, addition of G to PM exerted merely a slight stimulatory effect on respiration by a factor of 1.1 (1.0-1.2; Fig 2B), also observed in rat heart (1.2 [4]), horse *triceps brachii* (1.2 [42]), rabbit soleus (1.2 [26]), and human skeletal muscle (1.3 [43]). OXPHOS capacity (ADP-stimulated oxygen flux) was significantly higher with pyruvate&malate compared to glutamate&malate at physiological temperature (PM_*P*_ versus GM_*P*_; Fig 2B). This pattern was reversed at 4 °C, under which conditions N-OXPHOS capacity was higher with GM than PM (Fig 2C).

**Fig 2.**
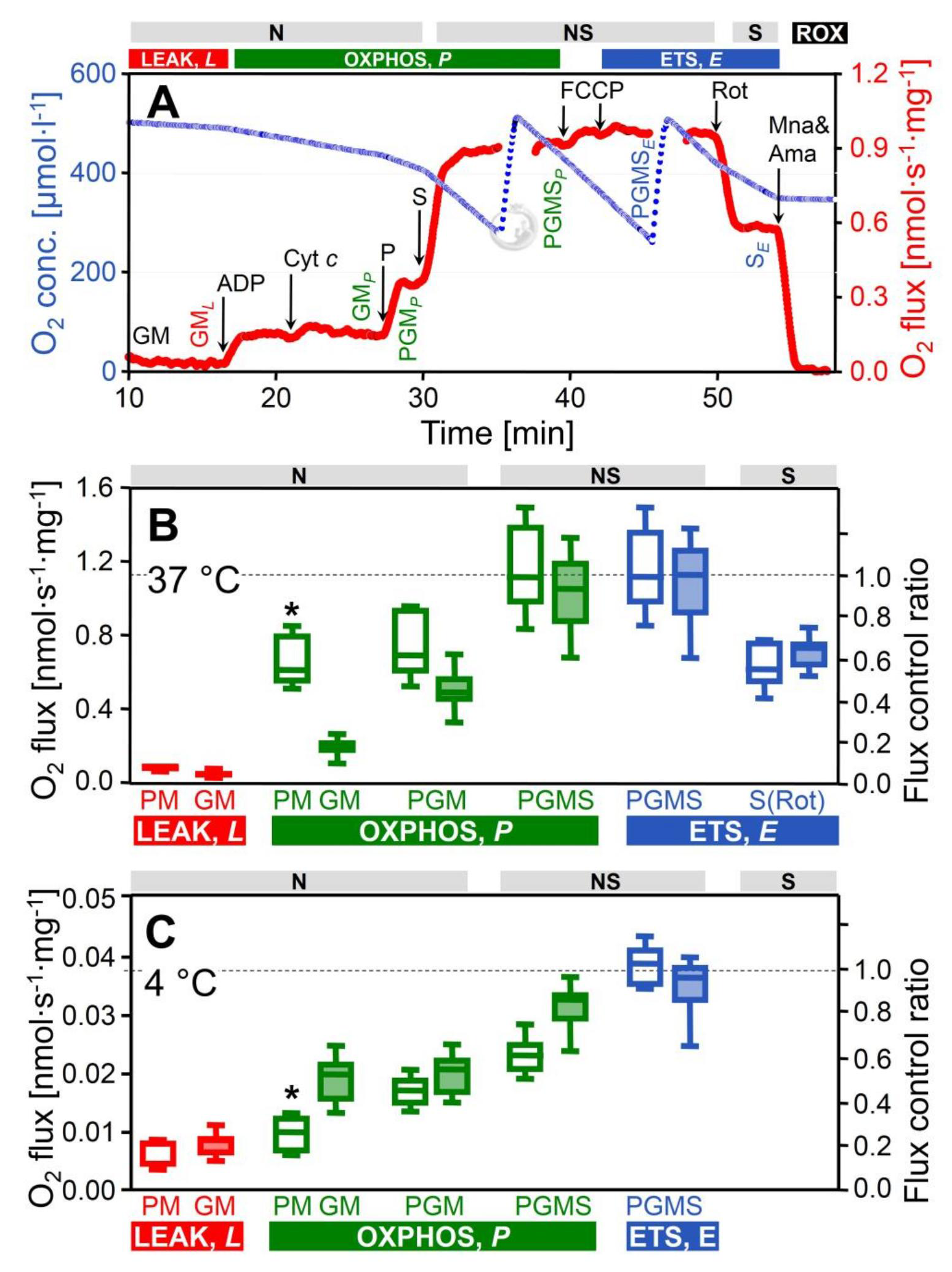
Respiration of permeabilized cardiac fibers (*J*_O2_, per wet weight), and flux control ratios normalized for NS-ETS capacity. (A) Oxygen consumption measured at 37 °C in SUIT 2 protocol: GM (N-LEAK respiration; GM_*L*_), ADP (N-OXPHOS capacity; GM_*P*_), cytochrome *c* (integrity of outer mt-membrane), pyruvate (N-OXPHOS capacity; PGM_*P*_), succinate (NS-OXPHOS capacity; PGMS_*P*_), FCCP (NS-ETS capacity; PGMS_*E*_), rotenone (Rot; inhibition of CI, S-ETS capacity; S_*E*_), malonic acid and antimycin A (Mna and Ama; less than 2% of residual oxygen consumption, ROX). (B, C) ROX-corrected respiration at 37 °C and 4 °C: LEAK respiration and OXPHOS capacity with PM (SUIT 1, open boxes, *n*=7 and 6 at 37 and 4 °C) or GM (SUIT 2, filled boxes, *n*=16 and 8 at 37 and 4 °C). Note the switch of theflux control pattern for N-OXPHOS capacity (PM_*P*_ vs. GM_*P*_). The two SUIT protocols merge at state PGM_*P*_, but results are shown separately (not significantly different). Flux control ratios are normalized relative to median PGMS_*E*_ of the combined protocols. Box plots indicate the minimum, 25^th^ percentile, median, 75^th^ percentile, and maximum. * Significant differences between the two SUIT protocols for the same state. For abbreviations see Fig 1.

In isolated mitochondria or permeabilized fibers incubated with PM, GM or PGM, citrate and 2-oxoglutarate are formed and rapidly exchanged for malate by the tricarboxylate and 2-oxoglutarate carriers, thus limiting the formation of succinate. In addition, succinate is lost into the incubation medium through the active dicarboxylate carrier exchanging succinate for inorganic phosphate. The high malate concentration equilibrates with fumarate, inducing product inhibition of succinate dehydrogenase (CII). Taken together, this limits succinate-linked electron transfer [44]. To simultaneously activate CII and NADH-related dehydrogenases of the TCA cycle, a high exogenous succinate concentration is required in addition to the NADH-linked substrates [45, 46], thus simulating the physiological condition of the NS-pathway with convergent electron flow into the Q-junction (Fig 1A). In the NS-pathway control state (PGMS_*P*_, Fig 1B), respiration almost doubled compared to OXPHOS capacity measured separately through the N-or S-pathway. The respiratory capacity through the S-pathway was similar to that observed with the NADH-linked substrate combinations. This provides evidence for an additive effect of convergent electron flow, expressed in terms of flux control ratios of N/NS = 0.53 (0.40-0.69), and S/NS = 0.61 (0.52-0.82) at 37 °C (Fig 2B). This additive effect was less pronounced at 4 °C (Fig 2C), but was similar between 25 °C and 40 °C (Fig 3).

**Fig 3.**
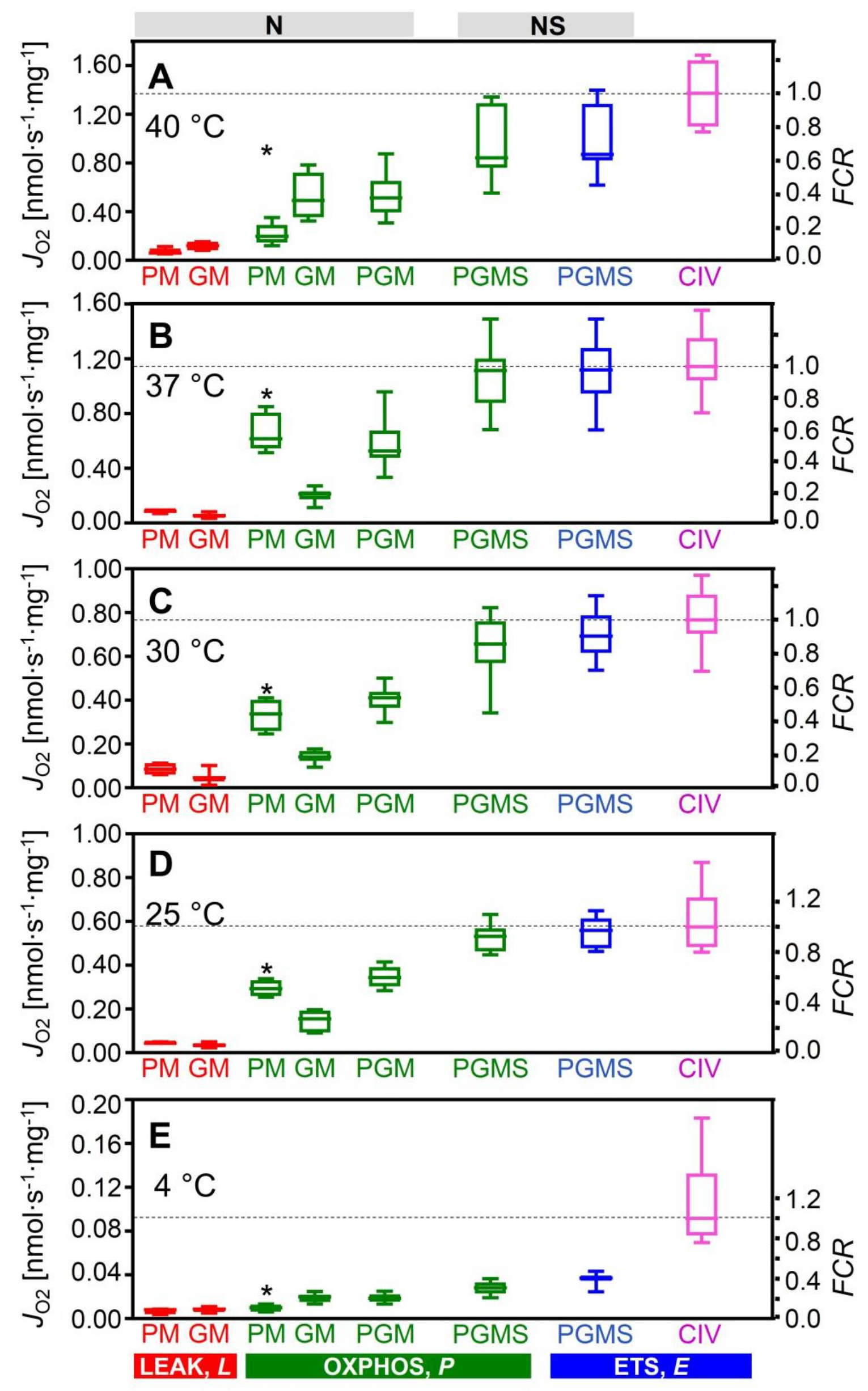
Respiration of permeabilized cardiac fibers (*J*_O2_, per wet weight), and flux control ratios, *FCR*, normalized for CIV activity at different temperatures. N_*L*_ and N_*P*_ with PM (SUIT 1; *n*=4-7) or GM(SUIT 2; *n*=5-16); N_*P*_ with PGM, NS_*P*_ and NS_*E*_ with PGMS (both protocols combined; *n*=9-23); normalized relative to median CIV_*E*_ (*n*=4-16). * Significant differences between the two SUIT protocols for the same state. For box plots and abbreviations see Fig 1 and 2.

### Temperature Sensitivity of Respiration Depends on Metabolic State

We determined temperature coefficients for evaluation of metabolic depression by hypothermia and specific remodeling of pathway control of OXPHOS (Table 1). Mitochondrial content, evaluated by the mitochondrial matrix marker citrate synthase activity, did not differ among tissue preparations. Citrate synthase activity [IU per mg fibers measured at 30 °C] was 0.249 (0.172-0.303), 0.234 (0.163-0.299), 0.236 (0.192-0.262), and 0.223 (0.184-0.279) in the 40, 37, 30 and 25 °C cohort, respectively. Therefore, divergences in mitochondrial content can be ruled out as a confounding factor. NS-OXPHOS capacity declined at 30 and 25 °C by 1.7- and 2.1-fold of the normothermic level (37 °C), and decreased slightly from 37 to 40 °C (Fig 3; Table 1). The thermal sensitivity of respiration was strongly dependent on coupling/pathway control states (Fig 4). The *Q*_10_ is the factor by which the reaction velocity increases for a rise in temperature of 10 °C. It varied between 1.2 and 2.4 across metabolic states in the temperature range of 25 to 37 °C (Fig 4C; Table 1). The *Q*_10_ for S-ETS capacity was 1.9 between 30 and 37 °C (Table 1), similar to results reported for heart mitochondria from guinea pigs (2.3) and rabbits (1.9) [32]. In this temperature range, *Q*_10_ values were close to 2.0 for NS-OXPHOS and ETS capacity, but an unexpectedly low *Q*_10_ of 1.4 was observed for PGM_*P*_. The N/NS flux control ratio (PGM/PGMS) increased with a decrease of temperature, from 0.53 (0.40-0.69) at 37 °C to 0.66 (0.58-0.72) at 25 °C, and remained high at 0.68 (0.55-0.79) at 4 °C (Fig 2C).

OXPHOS at 4 °C was only 3% of normothermic flux. This pronounced metabolic arrest resulted from the high thermal sensitivity of respiration with pyruvate (Table 1). *Q*_10_ for OXPHOS capacity with PM and PGMS increased to 4-5 under hypothermia from 25 to 4 °C. OXPHOS capacity with GM was much less temperature dependent, with a *Q*_10_ of 1.4 from 37 to 25 °C, and 2.7 from 25 to 4 °C. As a result, the OXPHOS capacity with PM, which was more than 2-fold higher than GM at physiological temperature, was depressed at 4 °C to a level even below GM-supported respiration (Fig 2C).

**Table 1.** Effect of temperature on mitochondrial respiration as a function of metabolic state in permeabilized mouse heart fibers. *Q*_10_ is the multiplication factor for an increase by 10 °C, calculated for respiratory flux, *J*, in the experimental temperature intervals, *T*_1_ to *T*_2_. Numbers in parentheses are multiplication factors (*MF* = *J*_*T2*_ / *J*_*T1*_) to convert *J* from *T*_1_ to *T*_2_. *Q*_10_ and *MF* are medians (*n*=4-16).

| State | | $Q_{10}$ (x $MF$) from $T_1$ to $T_2$ | | | |
| --- | --- | --- | --- | --- | --- |
|  |  | 4 °C to 25 °C | 25 °C to 37 °C | 30 °C to 37 °C | 37 °C to 40 °C |
| $CIV_{E0}$ | extrapolated | 1.84 (x 3.61) | 1.66 (x 1.83) | 2.17 (x 1.72) | 0.39 (x 0.76) |
| $CIV_E$ | measured | 2.40 (x 6.27) | 1.77 (x 1.99) | 1.77 (x 1.49) | 1.84 (x 1.20) |
| $NS_E$ | $PGMS_E$ | 3.64 (x 15.1) | 1.78 (x 2.00) | 1.98 (x 1.61) | 0.43 (x 0.78) |
| $NS_P$ | $PGMS_P$ | 4.06 (x 19.0) | 1.85 (x 2.10) | 2.13 (x 1.70) | 0.39 (x 0.76) |
| $S_E$ | $PGMS(Rot)_E$ | - | 1.93 (x 2.20) | 2.19 (x 1.73) | 0.25 (x 0.66) |
| $N_P$ | $PGM_P$ | 4.03 (x 18.7) | 1.42 (x 1.53) | 1.43 (x 1.28) | 0.91 (x 0.97) |
| $N_P$ | $PM_{CP}$ | 5.01 (x 29.5) | 1.86 (x 2.10) | 2.36 (x 1.82) | 0.48 (x 0.80) |
| $N_P$ | $PM_P$ | 4.24 (x 20.7) | 1.95 (x 2.22) | 2.39 (x 1.84) | 0.55 (x 0.84) |
| $N_P$ | $GM_{CP}$ | 2.67 (x 7.85) | 1.29 (x 1.36) | 1.79 (x 1.51) | 0.78 (x 0.93) |
| $N_P$ | $GM_P$ | 2.49 (x 6.81) | 1.39 (x 1.48) | 2.07 (x 1.67) | 0.65 (x 0.88) |
| $N_L$ | $PM_L$ | 2.23 (x 5.41) | 1.76 (x 1.97) | 1.01 (x 1.01) | 3.13 (x 1.41) |
| $N_L$ | $GM_L$ | 1.93 (x 3.99) | 1.45 (x 1.54) | 1.23 (x 1.16) | 1.38 (x 1.10) |

**Fig 4.**
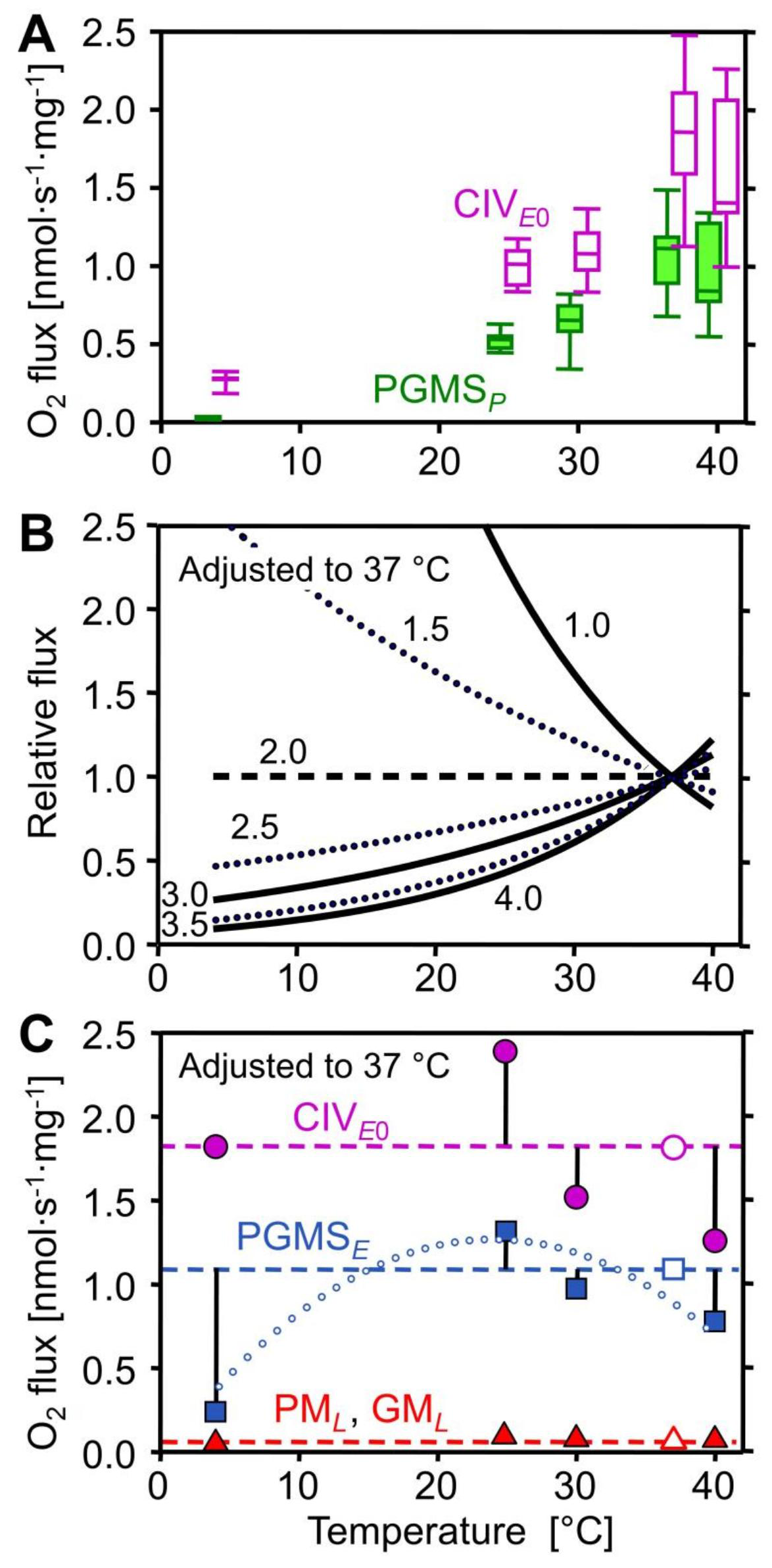
Effect of temperature on mitochondrial respiration. (**A**) Mass-specific oxygen flux (*J*_O2_) inrespiratory states PGMS_*P*_ (filled boxes) and CIV_*E*0_ extrapolated from the threshold plots in Fig. 5 (empty boxes). (**B**) Respiratory flux (*j*_O2_) relative to a simple temperature reference model: the reference flux isdefined at 37 °C as 1.0 and at other temperatures as 1.0 if *Q*_10_ is constant at 2.0 (horizontal dashed line). Non-linear deviations from standard conditions (full and dotted lines) are obtained when *Q*_10_ differs from 2.0 (*Q*_10_ shown by numbers) and is constant throughout the entire temperature range. **(C)** *j*_O2_ relative to standard temperature correction of flux at 37 °C (*Q*_10_ = 2.0; horizontal lines): CIV_*E*0_ (filled circles; extrapolated from the threshold plots, Fig. 5), ETS capacity (filled squares; PGMS_*E*_), and LEAK respiration (filled triangles; pooled GM_*L*_ and PM_*L*_). Vertical bars show deviations of experimental results (means, *n*=5 to 14) from the theoretical line for a *Q*_10_ of 2.0. The dotted trend line illustrates the change of temperature sensitivity for ETS, particularly at 4 °C. For box plots and abbreviations see Fig 1 and 2.

**Fig 5.**
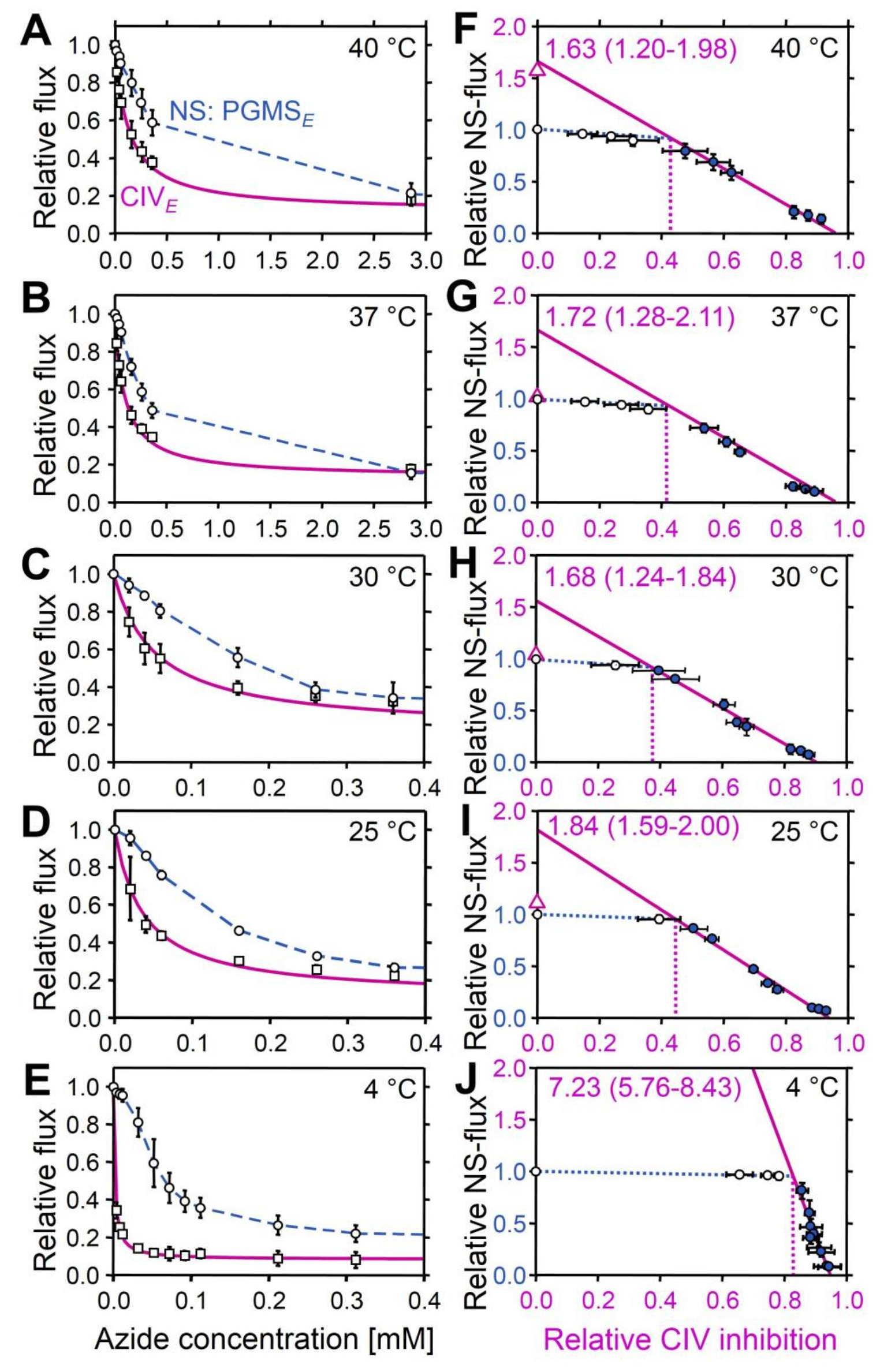
Azide titration and Complex IV threshold in permeabilized cardiac fibers at 40 to 4 °C. (**A** to**E**) Effect of azide titration on relative NS-pathway ETS capacity (PGMS_*E*_; circles, dashed line: linearinterpolations) and velocity of the single enzyme cytochrome *c* oxidase (CIV_*E*_; squares, solid line: hyperbolic fit). (**F** to **J**) Threshold plots of relative NS-pathway flux as a function of relative inhibition of CIV at identical azide concentrations. Data up to the threshold of inhibition are shown by open symbols. The CIV_*E*0_/NS_*E*_ flux ratio is calculated as the intercept at zero CIV inhibition of a linear regression through the data above the inflection point (closed symbols; *r* ^2^≥0.99). CIV_*E*0_/NS_*E*_ values are listed in the graphs as medians (min-max). The apparent excess capacity of CIV is *j*ExCIV = CIV_*E*0_/NS_*E*_ - 1. Triangles on *Y* axes show medians of relative CIV activities measured directly with TMPD and ascorbate (CIV_*E*_/NS_*E*_ =3.0 at 4 °C; not shown). The threshold of inhibition is at the intercept between the linear regression and the extrapolated line drawn from the control to the first inhibited flux (dotted vertical lines). Circles are means ± SD (*n*=4-5).

### Electron Transfer System Capacity and Coupling

Uncoupler titrations were performed in the ADP-activated state, to evaluate ETS capacity (*E*, noncoupled) in relation to OXPHOS capacity (*P*, coupled). *P* and *E* were numerically almost identical, indicating that the capacity of the phosphorylation system did not exert a limiting effect on respiration, with control located mainly at the level of the dehydrogenases (Fig 1A). Resting oxygen flux or LEAK respiration, *L*, is an inverse function of the proton/electron stoichiometry [47], and is therefore higher for the S-versus N-linked pathway. Respiration was measured in the presence of PM or GM before addition of ADP (N_*L*_) and after stimulation by ADP (N_*P*_). OXPHOS coupling efficiencies, 1-*L/P* [16], were independent of temperature in the range of 25 to 40 °C, and higher for PM than GM: 0.85 (0.78-0.89), 0.73 (0.57-0.80) and 0.82 (0.79-0.82) for PM, compared to 0.72 (0.54-0.81), 0.64 (0.39-0.89) and 0.69 (0.67-0.85) for GM at 37 °C, 30 °C, and 25 °C, respectively. At 4 °C, OXPHOS coupling efficiencies declined with GM to 0.58 (0.47-0.73), but even more with PM to 0.48 (0.28-0.60). The *Q*_10_ of LEAK respiration with GM and PM remained close to 2.0 down to 4 °C (Fig 4C; triangles).

### Apparent Cytochrome *c* Oxidase Excess Capacity

Since electron transfer capacity is limited when electrons are supplied only through the N-pathway, the major control resides under these conditions upstream in the dehydrogenases of the TCA cycle, with a correspondingly high apparent excess capacity of respiratory complexes downstream (for review see [13]). In physiological states with simultaneous NS-electron flow, the apparent excess capacity of downstream electron transfer is lower and flux control is shifted towards CIII and CIV. We therefore examined the apparent excess capacity of CIV at maximum convergent pathway flux through the ETS. Azide titrations resulted in hyperbolic inhibition of CIV (Fig 5A-E). The threshold plots display NS-pathway flux as a function of CIV activity (Fig 5F-J). The two distinct phases are related to (1) the elimination of excess capacity above the threshold (initial slope; dotted lines) and (2) the flux control coefficient below the threshold, where further inhibition of CIV causes a linear inhibition of pathway flux (full lines). Inhibition of CIV activity to 41% of controls exerted only a minor effect on respiratory capacity (37 °C; Fig 5G). The apparent excess capacity of CIV, *j*_ExCIV_ (for definition see Fig 5), at 37 °C was significant (median 0.72, range 0.28-1.11) with reference to convergent NS-electron flow, which provides the basis for a low flux control coefficient of CIV and a high functional threshold (Fig 5G). CIV excess capacities based on threshold plots were 0.6 to 0.8 at 40 to 25 °C (Figs 5F to I). The steep increase of *j*_ExCIV_ to 6.2 at 4 °C (Fig 5J) was associated with the high temperature sensitivity of ETS capacity upstream of CIV (Fig 3E).

## Discussion

Mitochondrial respiration in the living cell is supported by electron supply from multiple dehydrogenases and convergent electron entry into the Q-junction. Physiological respiratory capacity is underestimated in isolated mitochondria and permeabilized fibers when using simple substrate combinations such as PM supporting the NADH-linked pathway (N), or a single substrate for the succinate-linked pathway (S). N-OXPHOS capacity (PGM_*P*_=0.52 nmol O_2_·s^-1^·mg^-1^ wet weight) from the present study of permeabilized cardiac fibers from the mouse (Fig. 2B), is equal or slightly lower compared to *in vivo* maximal myocardial oxygen consumption (0.7 nmol O_2_·s^-1^·mg^-1^ wet weight [48]). However, PGM_*P*_ reached only 54% of NS-OXPHOS capacity (PGMS_*P*_=1.02 nmol O_2_·s^-1^·mg^-1^ wet weight), which actually exceeds maximal oxygen consumption of the working heart. Similarly, oxygen consumption of the perfused dog heart has been compared with N-OXPHOS capacity of isolated mitochondria expressed per tissue mass (0.42 and 0.50 nmol O_2_∙s^-1^∙mg^-1^ wet weight, respectively [49]). Under conditions of separate N-or S-pathway control, flux is limited artificially by selective substrate supply, thus effectively under-utilizing the apparent excess capacity of respiratory complexes downstream [16 and shifting flux control to dehydrogenases upstream

As expected on the basis of this respiratory flux control pattern, the apparent excess capacity of CIV with respect to N-OXPHOS capacity is high (2.2; unpublished observation), similar to apparent CIV excess capacities in permeabilized fibers from human skeletal muscle [51] and isolated mitochondria from rat heart and skeletal muscle (1.0 to 2.0 [52, 53]). Our data provide a rationale suggesting that these high apparent CIV excess capacities represent *in vitro* experimental artifacts which can be easily avoided. The apparent CIV excess capacity was lower when related to maximum ETS capacity of the NS-pathway (0.7 at 37 °C; Fig. 5), consistent with apparent CIV excess capacities in intact, uncoupled cells (0.0 to 0.4 [54–56]) and permeabilized human skeletal muscle fibers with NS-pathway control (0.4 [51]). Therefore, previously suggested discrepancies of CIV excess capacities in isolated mitochondria versus permeabilized fibers or intact cells [51, 54, 55] can be explained by NS-versus N-or S-pathway control states in mt-preparations and the concept of the Q-junction. Rocher al. [57] indicated that the quantity of mtDNA in human cell lines is tightly correlated to CIV activity. Importantly, apparent excess in catalytic capacity does not signify that it is not functionally required, considering the importance for high affinity of mitochondria to oxygen [52, 58] and the role of CIV in the control of cytochrome reduction levels [59].

Respiratory complexes form supercomplexes modulated by assembly factors, contributing to respiratory control and optimization of cellular metabolism [60]. Electrons are channeled through supercomplexes (CI+III [61] and CI+III+IV [62] in the mouse heart) which restrict exchange with the free Q- and cytochrome *c*-pools, thus limiting random collisions [61, 63, 64]. Under conditions of tight electron channeling, activation of an additional convergent pathway would exert a completely additive effect on ETS capacity. Our results provide evidence against maximally tight supercomplex channeling in the mouse heart, since the combined NS-pathway capacity was significantly less than predicted from a completely additive effect of the N-and S-pathway fluxes measured separately. Two other mechanisms could reduce the additive effect on ETS capacity: (i) substrate comp], and (ii) limitation of electron transfer by deficient enzyme capacities downstream of the Q-cycle in the absence of tight channeling. The additive effect on flux of convergent electron transfer at the Q-junction varies between species, strains, tissues, age, and pathophysiological conditions, unraveling an unexpected diversity of mitochondrial respiratory control patterns [25].

The phosphorylation system represents another functional unit potentially contributing to the limitation of OXPHOS capacity, *P*, relative to ETS capacity, *E*. The extent of this limitation, *E-P*, is expressed by the excess *E-P* capacity factor*, j_ExP_* = (*E-P*)/*E* = 1-*P/E*, ranging from zero (no limitation) to the upper limit of 1.0 [16]. *j_ExP_* was very low but significantly different from zero in mouse heart (0.03 at 25, 37, and 40 °C, and 0.05 at 30 °C; *P*<0.05, pooled protocols; Fig 3), while it is zero in mouse skeletal muscle [24] and rat heart [20], and 0.04 and 0.02 in rat soleus and extensor digitorum longus muscle, respectively [23]. In human skeletal muscle, however, the limitation by the phosphorylation system is highly significant, *j_ExP_*(NS) = 0.05 to 0.2 [6, 65] and *j_ExP_*(N) = 0.2 with GM as substrates [66], and is even more pronounced in the human heart, with *j_ExP_*(N) increasing from 0.52 in healthy controls to 0.59 in heart failure [2].

Temperature exerts complex effects on coupling control of mitochondrial respiration [67]. Proton leak is a property of the inner mt-membrane and depends on mt-membrane potential, whereas proton slip is a property of the proton pumps and depends on enzyme turnover. Fierce controversies on proton leak versus slip, which collectively control LEAK respiration, could be resolved by considering physiological (37 °C) versus conventional ‘bioenergetic’ temperature (25 °C) [67]. Temperature coefficients vary between different enzyme-catalyzed reactions involved in mitochondrial respiration (Table 1). Relatively small differences of *Q*_10_ among the different enzymes of a pathway may have dramatic impact on metabolic organization and OXPHOS remodeling over a wide range of temperatures. The excess *E-P* capacity factor in the mouse heart increased at 4 °C to 0.36 (0.33-0.51) and 0.09 (0.03-0.16) in SUIT 1 and 2, respectively (Fig 2C), consistent with an increase of the flux control coefficient of the phosphorylation system at low temperature in rat liver mitochondria [67, 68]. Furthermore, the apparent CIV excess capacity in mouse heart increased strikingly at 4 °C compared to normothermic values (Fig. 5). The excess of CIV relative to NS-pathway capacity increases in peritoneal but not in alveolar murine macrophage-derived cell lines at 25 °C compared to 37 °C [69]. Although a decline of CI activity with temperature is possibly involved in the shift of CIV excess capacity [69], our results suggest a different mechanism. The highest OXPHOS *Q*_10_ of 5.0 was obtained with pyruvate&malate between 4 and 25 °C, in contrast to the lower *Q*_10_ with glutamate&malate (Table 1). Both substrate combinations fuel the N-pathway with electron entry into the Q-junction through CI. Hence CI activity cannot explain the observed hypothermic response pattern. This confirms a previous study in rat heart mitochondria showing a high thermal sensitivity of pyruvate-supported respiration and activity of the PDC complex at low temperature [70].

The thermal sensitivity of PDC or pyruvate supported respiration was not as pronounced in a cold adapted fish species (*Anarhicas lupus*) [71], suggesting that pyruvate dehydrogenase plays not only an important role in the response of OXPHOS to temperature in the murine heart [70], but is also a potential site of key adaptation to upgrade mitochondrial capacity at low temperature in ectotherm species [72]. This is in line with evidence of control of convergent OXPHOS at upstream steps of electron supply in *Fundulus heteroclitus* heart mitochondria [73, 74]. A higher control of the phosphorylation system is observed at lower temperature (17 °C compared to 25 °C) in permeabilized heart fibers from the triplefin fish (*Forsterygion lapillum*) [75]. N-or S-linked respiration appeared to be more affected by temperature changes at lower temperature compared to CIV in the freshwater turtle *Trachemys scripa* [76]. Similarly, permeabilized muscle fibers from *Drosophila simulans* show an increase of CIV excess capacity at low temperature [77]. Taken together, NADH-linked mt-matrix dehydrogenases and the phosphorylation system rather than electron transfer complexes appear to be the primary modulators of respiratory control patterns at low temperature in mitochondria from endotherm and ectotherm species.

The conventional respiratory acceptor control ratio, RCR (State 3/State 4 [17] or *P/L* ratio) was 6.7 with PM (37 °C), and much lower (3.6) with GM. For statistical and conceptual reasons, the RCR is replaced by the OXPHOS coupling efficiency, 1-*L/P* = 1-RCR^-1^ [16]. The low OXPHOS coupling efficiency for GM of 0.72 (compared to 0.85 for PM) reflects the low OXPHOS capacity supported by specific substrates, rather than low coupling efficiency. Evaluation of coupling must be based on pathway control states supporting a high ETS capacity under conditions not limited by substrates or by the phosphorylation system. Hence, PM as a 'fast' pathway at 37 °C yields a more appropriate index of coupling. Under deep hypothermia (4 °C), however, the estimation of coupling is complicated by the strong thermal sensitivity of ETS capacity compared to LEAK respiration (Fig 3B) and by the limitation imposed by the phosphorylation system on OXPHOS capacity.

Our results show that the estimation of maximal respiratory capacity should be performed at physiological temperature and with substrate combinations appropriate to ensure suitable operation of the TCA cycle and multiple entries of electrons into the Q-cycle. Those conditions define the physiological reference state for mitochondrial respiratory control. This is of major significance to better explain the pathological effects of genetic mutations and acquired CIV deficiencies. Furthermore, our results on the key modulators of thermal sensitivity of mitochondrial metabolism will allow to determine how cells and tissues are impaired by temperature changes, including modest and deep hypothermia used for cardiac surgery. This novel aspect of mitochondrial medicine may provide a basis for intervention strategies to limit damages [69]. From an evolutionary perspective, our study allows to pinpoint candidate loci that are potentially under selective pressure and therefore represent targets for seeking mechanisms of adaptation to different temperature regimes. In summary, we propose the hypothesis that restitution of the loss of respiratory capacity at low temperature requires compensatory adaptations to counteract upstream limitations of ETS capacity at the entry to and within the TCA cycle (particularly the PDC) and downstream limitation of flux by the phosphorylation system.

## Materials and Methods

### Preparation of Permeabilized Myocardial Fibers

Adult male mice C57 BL/6N were housed under standard conditions according to the Austrian Animal Care Law. At 8 to 10 weeks of age (23 ± 3 g), animals were anaesthetized with ketamine and xylazine (80 and 10 mg∙kg^-1^, respectively) given intramuscularly. The heart was excised and placed in 5 ml of ice-cold relaxing solution BIOPS [6]. After rapid mechanical permeabilization of the left ventricle (~40 mg wet weight), bundles of fibers were agitated gently (30 min, 4 °C) in BIOPS supplemented with 50 μg∙ml^-1^ saponin [78]. Fibers were washed by agitation (10 min, 4 °C) in mitochondrial respiration medium MiR05 [40], immediately blotted, weighed, and used for respirometric measurements.

### High-Resolution Respirometry

Respiration was measured simultaneously in 10 respiration chambers (Oroboros Oxygraph-2k, Innsbruck, Austria), one Oxygraph-2k with two chambers for each of the following temperatures: 4, 25, 30, 37, and 40 °C. Permeabilized fibers (0.7-1.3 mg at 25 to 40 °C and 7-8 mg at 4 °C) were used in each chamber containing 2 ml of MiR05. Respiratory flux was expressed per mg wet weight of fibers. Instrumental and chemical oxygen background fluxes were calibrated as a function of oxygen concentration and subtracted from the total volume-specific oxygen flux ([52, 79]; Datlab software, Oroboros Instruments). An oxygen regime of 500 to >200 µM was maintained at 30 to 40 °C, but up to 700 and 900 µM at 25 and 4 °C, to avoid artificial oxygen diffusion limitation of flux [80]. In the first substrate-uncoupler-inhibitor titration (SUIT) protocol, the following final concentrations were added sequentially: G (10 mM), M (5 mM), P (5 mM), ADP (1 mM), cytochrome *c* (10 μM), S (10 mM), FCCP (optimum concentration, 0.125 to 0.375 μM), Rot (0.5 μM), Ama (2.5 μM), Mna (5 mM), TMPD (2 mM) and ascorbate (0.5 mM). In the second SUIT protocol addition of G and P were inversed. An increase of respiration due to cytochrome *c* addition after ADP was observed at 30 to 40 °C, with cytochrome *c* control factors (change of respiration divided by cytochrome *c* stimulated respiration [16]) in the range of 0.05 to 0.15, with higher values of 0.11 to 0.20 at 25 °C. At 4 °C, N-OXPHOS capacity showed a trend to decline during the experiment particularly with PM, and no stimulation could be observed with ytochrome *c*. Thus the integrity of the outer mitochondrial membrane in mouse heart permeabilized fibers was comparable to rat heart fibers studied at 30 °C [81]. Residual oxygen consumption (ROX), evaluated after inhibition of CI, CII and CIII with rotenone, malonic acid and antimycin A (Rot, Mna and Ama, respectively) was a small fraction (0.01 to 0.02) of NS-linked ETS capacity at 25 to 40 °C, but increased to 0.04 to 0.10 at 4 °C. Nevertheless, correction of fluxes in all respiratory states for ROX was significant, particularly in the resting state of LEAK respiration, when ROX was as high as 0.12 to 0.32 of total oxygen consumption in the N-linked LEAK state at 25 to 40 °C.

Apparent CIV excess capacities were determined by azide titrations of CIV activity and of NS-ETS capacity at 4, 25, 30, 37, and 40 °C. Threshold plots of relative respiration rate against the fraction of inhibited CIV activity at the same azide concentration were made as previously described [54, 82]. Azide titrations were performed at optimum uncoupler concentration supporting maximum flux, preventing the effect of inhibition of ATP synthase [83] and eliminating any contribution of the phosphorylation system to flux control. The following azide concentrations were used (in mM): 0.02, 0.04, 0.06, 0.16, 0.26, 0.36, 2.9, 5.4, 10.4 between 25 and 40 °C, and 0.004, 0.008, 0.012, 0.032, 0.052, 0.072, 0.092, 0.11, 0.21, 0.31, 2.8, 5.3, 10.3 at 4 °C (not all points shown in Fig 5 due to overlap).

The contents of the chambers were removed at the end of each experimental run and the chamber was rinsed twice with 500 μl of respiration medium. The fibers were homogenized for 2 × 30 s with an Ultra-Turrax homogenizer at maximum speed and immediately frozen in liquid nitrogen and stored at -80 °C for subsequent measurement of citrate synthase at 30 °C [84].

### Data Analysis

Statistica software^©^ was used for statistical analyses. Data were log transformed to meet the requirement for heteroscedasticity according to Levene’s test. A three-factor ANOVA (protocol, state and temperature) followed by *a posteriori* Tukey multiple comparison tests were used to test for differences between protocols at a specific state and temperature. To determine the effects of addition of substrates, cytochrome *c*, inhibitors, or uncoupler, a t-test for dependent samples was used. Significance was considered at *P*<0.05. Results are presented without transformation as medians (min-max) unless specified otherwise.

## Acknowledgments

We acknowledge the support by J-C Tardif and R Margreiter. M Schneider performed CS analysis. We thank CL Hoppel and B Tandler for helpful comments.

## Author contributions

Conceptualization: Erich Gnaiger, Hélène Lemieux, Pierre U. Blier.

Formal analysis: Erich Gnaiger, Hélène Lemieux.

Funding acquisition: Erich Gnaiger, Pierre U. Blier, Hélène Lemieux.

Investigation: Hélène Lemieux, Erich Gnaiger.

Methodology: Hélène Lemieux, Erich Gnaiger.

Project admninistration: Erich Gnaiger.

Ressources: Erich Gnaiger.

Supervision: Erich Gnaiger.

Visualization: Hélène Lemieux, Erich Gnaiger.

Writing – Review & Editing: Hélène Lemieux, Pierre U. Blier, Erich Gnaiger.

## Funding sources

HL received scholarships from the Canadian NSERC and the Quebec FQRNT. Funding for this work was provided by an NSERC discovery grant to PUB and HL (RGPIN 155926 and 402636), K-Regio project MitoFit (EG) and Oroboros Instruments. Contribution to COST Action CA15203 MITOEAGLE.

